# Flow cytometry methods for targeted isolation of ctenophore cells

**DOI:** 10.1101/2023.08.10.552885

**Authors:** Abigail C Dieter, Aliyah BK True, Emily Gilbertson, Grace Snyder, Adam Lacy-Hulbert, Nikki Traylor-Knowles, William E Browne, Lauren E Vandepas

**Author notes:** **Corresponding Author:** Lauren E Vandepas. Authors contributed equally to this work (Abigail C Dieter, Lauren E Vandepas).

## Abstract

Cell suspension fluidics, such as flow cytometry (FCS) and fluorescence-activated cell sorting (FACS), facilitates the identification and precise separation of individual cells based on phenotype. Since its introduction, flow cytometry has been used to analyze cell types and cellular processes in diverse non-vertebrate taxa, including cnidarians, molluscs, and arthropods. Ctenophores, which diverged very early from the metazoan stem lineage, have emerged as an informative clade for the study of metazoan cell type evolution. We present standardized methodologies for flow cytometry-mediated identification and analyses of cells from the model ctenophore *Mnemiopsis leidyi* that can also be applied to isolate targeted cell populations. Here we focus on the identification and isolation of ctenophore phagocytes. Implementing flow cytometry methods in ctenophores allows for fine scale analyses of fundamental cellular processes conserved broadly across animals, as well as potentially revealing novel cellular phenotypes and behaviors restricted to the ctenophore lineage.

## 1 Introduction

Flow cytometry (FCS) was developed as a technique to analyze intrinsic cellular properties of mammalian cells, including relative cell size and presence of intracellular granules (Fulwyler, 1965; Herzenberg et al., 2002). Light scatter properties and fluorescence, measured by lasers and photon emission detectors, are used to characterize parameters of individual cell morphologies as well as a wide range of molecular labels. Thus, flow cytometry represents a powerful tool for identifying and investigating individual cells within a heterogeneous cellular suspension (Barteneva et al., 2012). Basic attributes associated with an individual cell, such as relative size and granularity (or complexity), can be measured via light scatter properties. Forward scatter (FSC) is used as a proxy for evaluating relative cell size. Side scatter (SSC), the measure of light scattered 90° from the source, is used as a proxy for determining the relative granularity of a cell (McKinnon, 2018). Fluorescence-activated cell sorting (FACS) further facilitates the identification of single cells based on the detection of subcellular fluorescent markers associated with specific cellular characteristics, such as the expression of specific proteins, cell cycle state, cell proliferation, cell viability, and apoptosis (Julius et al., 1972; Adan et al., 2017; McKinnon, 2018). FACS has also been used successfully to analyze cellular processes in diverse non-vertebrate marine organisms including corals, tunicates, and molluscs (de la Cruz et al., 2008; Choi et al., 2010; Schippers et al., 2011; Park et al., 2012; Rosental et al., 2017; Cheng et al., 2018; Siebert et al., 2019; Snyder et al., 2020).

Ctenophora are a clade of non-bilaterian, gelatinous marine predators possessing a suite of unique character traits (Dunn et al., 2015). Genomic analyses place Ctenophora near the base of the Metazoa, thus making them a critical group to study evolution of metazoan cell types (Sebé-Pedrós et al., 2018; Li et al., 2021; Schultz et al., 2023). Previous studies have usually relied upon microscopy-based analyses to characterize distinct ctenophore cell types including true muscle cells, nerve cells, various digestive cells, stellate phagocytic amoebocytes, and ctenophore-specific cell types including tentacular colloblasts and ctene-row ciliary cells (comb-rows) (Hernandez-Nicaise, 1991; Jager et al., 2011; Dayraud et al., 2012; Moroz et al., 2014; Presnell et al., 2016; Babonis et al., 2018; Traylor-Knowles et al 2019; Jokura et al., 2022; Burkhardt et al., 2023). Additionally, optimization of methodologies that build on cell culture techniques (Presnell et al., 2016; Vandepas et al., 2017; Dieter et al., 2022) are required to study functional characteristics and traits associated with specific cell types. Here we present reliable, repeatable methods for performing flow cytometry and FACS with ctenophore primary cells that facilitate the isolation, behavioral assessment, and functional characterization of distinct cell types in the model lobate ctenophore *Mnemiopsis leidyi*. We apply these methods for the identification, isolation, and collection of ctenophore phagocytes.

## 2 Materials and Methods

### 2.1 Animal maintenance and preparation of ctenophore cells

#### 2.1.1 Laboratory culture of Mnemiopsis

Laboratory strains of *Mnemiopsis leidyi* were maintained as previously described (Presnell et al., 2022). Individual animals were isolated in minimal ctenophore media (MCM); 0.22 μm filter-sterilized filtered seawater (FSW; Instant Ocean) treated with 1x penicillin/streptomycin solution (P/S) (100 units/mL penicillin, 100 μg/mL streptomycin; Gibco; ThermoFisher). Isolated animals were deprived of food for a minimum of three hours and subsequently screened via light microscope to verify clearance of gut contents. Three water changes with FSW were then performed to remove remaining debris and excess mucus.

#### 2.1.2 Dissociation of ctenophore tissues and cell isolation

Ctenophore cells were mechanically dissociated using a dounce homogenizer and loose fitting pestle as previously described (Vandepas et al., 2017; Dieter et al., 2022). Briefly, individual adult ctenophores were transferred into FSW supplemented with 2x P/S immediately prior to homogenization with 10-15 strokes of the loose fitting pestle. The homogenization step should be performed slowly to reduce shear forces that may damage cells, leading to poor viability and/or yield.

Dissociated cells were decanted to a 15 mL centrifuge tube (Falcon). An equal volume of chilled FACS buffer (0.2 μm filter-sterilized 1x PBS, 2%(v/v) FBE (fetal bovine essence, Avantor),1% penicillin (Sigma, P7794), 1% streptomycin (Sigma, S9137), 2 mM EDTA (Sigma, E5134), and 400 mM NaCl (Sigma, S3014); stored at 4°C) was then gently mixed into the cell homogenate by inversion. The cell suspension was centrifuged at 500 *g* for 10 minutes at 8°C to pellet the cells. The upper, cell free, supernatant was then removed by aspiration, leaving behind a translucent, loosely pelleted visible cell fraction. An additional wash was performed by gently resuspending the cell pellet with approximately 2x volume of chilled FACS buffer and re-filtering the cell homogenate through a cell strainer stack with a final 70 μm mesh (pluriStrainer) to remove any remaining large cell aggregates and/or tissue debris (Dieter et al., 2022). Cell suspension densities were determined using an automated cell counter (Invitrogen Countess 3 FL) and adjusted to ∼1-2 ×10^6^ cells/mL with chilled FACS buffer. Prepared cell suspensions are kept on ice prior to flow cytometry analysis.

While the methods and representative results presented here use cell preparations isolated from individual ctenophores diluted to ∼1-2 ×10^6^ cells/mL, applications requiring large numbers of cells (for example, single-cell sequencing) can combine multiple individuals. However, increasing relative cell densities can result in increased viscosity of the cell suspension, which may reduce flow or clog microfluidic chambers. The addition of one or more filtration steps with a 70 μm mesh prior to cell sorting can mitigate reductions in flow within microfluidic chambers when analyzing high density cell preparations.

#### 2.1.3 Assessing viability of Mnemiopsis cell preparations

To determine cell viability, readily available reagents used to determine mammalian cell culture viability were compared: Trypan Blue (Invitrogen 15250061) and ReadyCount Green/Red Viability Stain (Invitrogen A49905) (Dieter et al., 2022). Both of these commercial stain preparations were mixed at a 1:7 ratio with ctenophore cell suspensions to mitigate osmotic differences between mammalian and ctenophore cell culture media. Quantification of cell viability was performed with an automated cell counter (Invitrogen Countess 3 FL). Cell suspensions with < 85% viability were not used for downstream flow cytometry assays.

### 2.2 Fluorescent Markers

To analyze samples containing fluorescent reagents, unstained controls were used in all experiments to visualize shifts in recorded event fluorescence and inform gating strategies.

#### 2.2.1 Propidium Iodide (eScience, USA)

Propidium iodide (PI) is a membrane impermeable DNA dye commonly used to determine cell viability (Riccardi and Nicoletti, 2006). Fluorescent PI signal correlates with nuclear staining of dead or dying cells that have compromised cell membranes (Johnson et al., 2013). *Mnemiopsis* cell suspensions were incubated with 2 μg/mL PI solution in MCM at room temperature for 15 minutes protected from light. After PI incubation, cell suspensions were centrifuged at 500 *g* for 10 minutes at 8°C to pellet cells. The supernatant was removed by aspiration. Cell pellets were gently resuspended with approximately 2x pellet volume of chilled FACS buffer and then filtered through a 70 μm mesh cell strainer.

#### 2.2.2 Vybrant DyeCycle Green (Invitrogen V35004, USA)

Vybrant DyeCycle Green is a cell permeable dye that exhibits stoichiometric binding with double stranded DNA. Fluorescent intensity of Vybrant DyeCycle Green increases linearly with DNA content, making it an efficient reagent for detecting cell cycle state (Kim and Sederstrom, 2016). Vybrant DyeCycle Green was used at a final concentration of 5 μM in MCM. *Mnemiopsis* cell suspensions were incubated at 16°C for 1 hr protected from light. After incubation, cell suspensions were centrifuged at 500 *g* for 10 minutes at 8°C to pellet cells. The cell free supernatant was then removed by aspiration. Cell pellets were gently resuspended with approximately 2x pellet volume of chilled FACS buffer and filtered through a 70 μm mesh cell strainer.

### 2.3 Preparation of *Mnemiopsis* Cells for Phagocytosis Assays

To functionally probe the phagocytic potential of ctenophore cells, we exposed heterogenous *Mnemiopsis* cell suspensions to pHrodo Red *E. coli* BioParticles (Invitrogen P35361). Fluorescence of labeled *E. coli* is selectively activated when exposed to low pH environments, such as within the lumen of phagosomes, endosomes or lysosomes (Kissing *et al*., 2018).

#### 2.3.1 Preparation of pHrodo E. coli BioParticles

Stock *E. coli* BioParticles were resuspended at 2 mg/mL in MCM. To disperse *E. coli* aggregates, the suspension was passed through a 28g needle fitted with a 1mL syringe 20 times. Alternatively, *E. coli* BioParticle suspensions can be disaggregated with sonication for 10 minutes.

#### 2.3.2 Preparation of Mnemiopsis cells for incubation with E.coli Bioparticles

*Mnemiopsis* cell suspensions were prepared as described in Section 2.1.2. Pelleted cells were then resuspended in MCM with pHrodo Red *E. coli* BioParticles at a final concentration of 100 μg/mL and incubated at RT for 1 hour protected from light on a gentle rocking platform to prevent cell suspensions from settling. The pHrodo *E. coli* BioParticles reach maximum fluorescence approximately 90 minutes after ingestion. After incubation, cell suspensions were centrifuged at 500 *g* at 8°C for 10 minutes. The upper cell free supernatant was decanted or aspirated to remove excess non-phagocytosed *E. coli*. The recovered cell pellet was then resuspended in 1mL chilled FACS buffer and filtered through a 70 μm Nylon mesh cell screen (Corning, cat# 431751), or alternatively a 70 μm pluriStrainer fitted with a Luer-Lock adapter ring and syringe over a 50mL Falcon tube (Dieter et al., 2022).

### 2.4 Flow cytometry

#### 2.4.1 Overview of Flow Cytometry Analysis

Flow cytometry assays were conducted using both a Sony SH800 Cell Sorter (LE-Sony SH800 V2.1.6) and BD FACSAria™ Fusion Flow Cytometer (BD FACSDiva™ Software) to compare FSC-A/SSC-A measurements and clustering sensitivity (Figure 1). Data was collected using LE-Sony SH800 V2.1.6 software and BD FACSDiva™ Software, respectively, and subsequently analyzed with FlowJo™ Software for Windows Version 10 (Becton, Dickinson and Company, Ashland, OR, USA). Flow cytometer nozzle size should be selected and optimized based on considerations for each assay and the size of targeted cell populations to reduce <u>s</u>orter-induced <u>c</u>ell <u>s</u>tress (SICS). For collecting a range of ctenophore cells, including phagocytes, 100 μm sorting chips (Sony Biotechnology Inc, San Jose, California, USA) were used. In addition, selection of the cytometer flow rate (sample pressure) for live cell sorting should be optimized to reduce cell damage. For collecting live ctenophore cells, the flow rate was set to 3. Typically, large nozzles (100 μm) and low flow pressure should be applied for sampling that will include the collection of large cells. A minimum of 30,000 cells were analyzed per sample.

**Figure 1:**
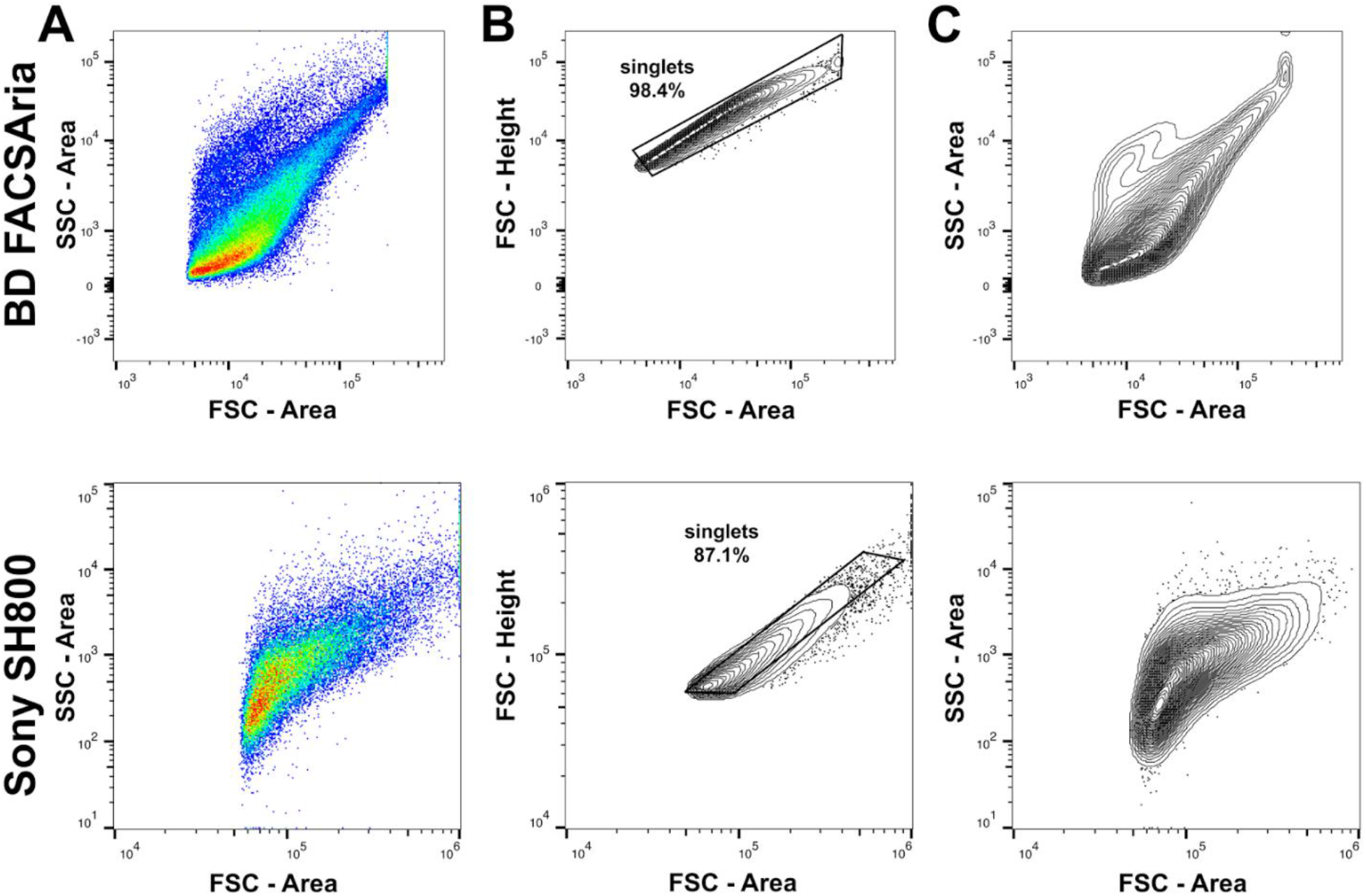
Comparisons of flow cytometry analyses of *Mnemiopsis* cells using a BD FACSAria™ Fusion Flow Cytometer (top row) and Sony SH800 Cell Sorter (bottom row). Both flow cytometers detected *Mnemiopsis* cells of variable size and granularity, as anticipated when querying a sample containing mixed cell types. A) FSC-A/SSC-A profiles of *Mnemiopsis* cell suspensions prior to doublet exclusion. B) Gating strategy for doublet and multiplet exclusion using FSC-H/FSC-A. C) Contour plot (2%) showing event densities of FSC-A/SSC-A profiles following doublet exclusion with outliers included.

Doublet and multiplet events that represent cell aggregates were removed and excluded from further analysis by selecting a diagonal gate around events with an approximate 1:1 ratio between FSC-Area and FSC-Height (Figure 1B). After applying this exclusion gate, the majority of recorded events represent single cells (Figure 1C). As expected, both flow cytometers detect *Mnemiopsis* cells of variable size and granularity when querying a sample containing mixed cell types (Figure 1C).

#### 2.4.2 Sony SH800 Sheath Buffer for Marine Invertebrate Cell Cultures

Sheath fluid is used to hydrodynamically focus cells in suspension as they travel through the Sony SH800 cytometer. We use a modified high-salt sheath buffer composed of 3X PBS (for 1L: 700mL ultrapure water, 300mL sterile 10X PBS) to reduce osmotic differences between the sheath fluid and the FACS buffer used to resuspend marine invertebrate cell preparations. Changes to the standard sheath buffer were not required for analyses of *Mnemiopsis* cells using a BD FACSAriaII Fusion flow cytometer, as the microfluidics for that system reduce mixing between the cell suspensions in FACS buffer and the sheath fluid.

#### 2.4.3 Sony SH800 Machine Maintenance Following Processing of High Salinity Samples

Several adjustments to standard cytometer maintenance were necessary to analyze cell preparations when using a high-salt sheath buffer on the Sony SH800. Prior to analyses, the flow cytometer collection chamber was washed with 70% ethanol and then cleaned with low-lint paper wipes to reduce triboelectric effects from static electricity build-up during machine operation. To prevent salt accumulation, both the waste collection chamber and deflection plates were periodically removed, soaked in ultrapure water and cleaned with 70% ethanol to remove remaining residual water from the cytometer components.

### 2.5 Imaging

Post sorting, cells were collected into 1.5 mL microcentrifuge tubes containing 500 μL of MCM. The collected cells were pelleted by centrifugation at 500 *g* at 16°C for 10 minutes. The cell-free supernatant was carefully removed, leaving the pellet of sorted cells in approximately 50 μL of MCM. The cell pellet was then gently resuspended using a sheared pipette tip. For microscopy, resuspended cells were pipetted onto microscope slides in 7.5 μL aliquots (Fisherbrand™ Superfrost™ Plus, VWR: 48311-703; fitted with SecureSeal Imaging Spacers, Electron Microscopy Sciences #70327-9S). Differential interference contrast (DIC) and fluorescent images were acquired using a Zeiss Axio Imager.Z2, Zeiss AxioCam MRm Rev3 camera and Zeiss Zen Blue software.

## 3 Results

### 3.1 Viability of *Mnemiopsis* Cell Suspensions

Identification and removal of dead or dying cells present in a cell suspension from subsequent flow cytometry analyses is a critical step to ensure accuracy of results. Additionally, measuring cell viability can be informative when assaying cellular responses to drugs or other experimental treatments (Kummrow et al., 2013). To quantify viability of *Mnemiopsis* cell preparations prior to flow cytometry analyses, live/dead cell counts were performed using Trypan Blue or ReadyCount Green/Red Viability Stain and an Invitrogen Countess Cell Counter. Across cell suspension preparation replicates, the Trypan Blue exclusion assay indicated an average of 7% cell death and the ReadyCount Green/Red Viability Stain indicated approximately 11% cell death (Table 1). We also incubated cells with propidium iodide (PI) and visualized PI fluorescence using FACS to independently assess *Mnemiopsis* cell preparation viability (Figure 2A-B). An increase in PI fluorescence compared to unstained controls identifies dead or dying cells (Figure 2B). PI staining indicated an average of 9% cell death across cell preparation replicates (Supplemental Fig. 1).

**Table 1:** Percent viabilities of *Mnemiopsis* cell suspension replicates using common staining assays.

| Biological Replicate | Trypan Blue Exclusion Assay | ReadyCount Viability Stain | Propidium Iodide (Flow cytometry) |
| --- | --- | --- | --- |
| Ctenophore 1 | 91 | 92 | 90 |
| Ctenophore 2 | 94 | 85 | 93 |
| Ctenophore 3 | 91 | 92 | 91 |
| Ctenophore 4 | 92 | 83 | 92 |
| Ctenophore 5 | 99 | 93 | 88 |

**Figure 2:**
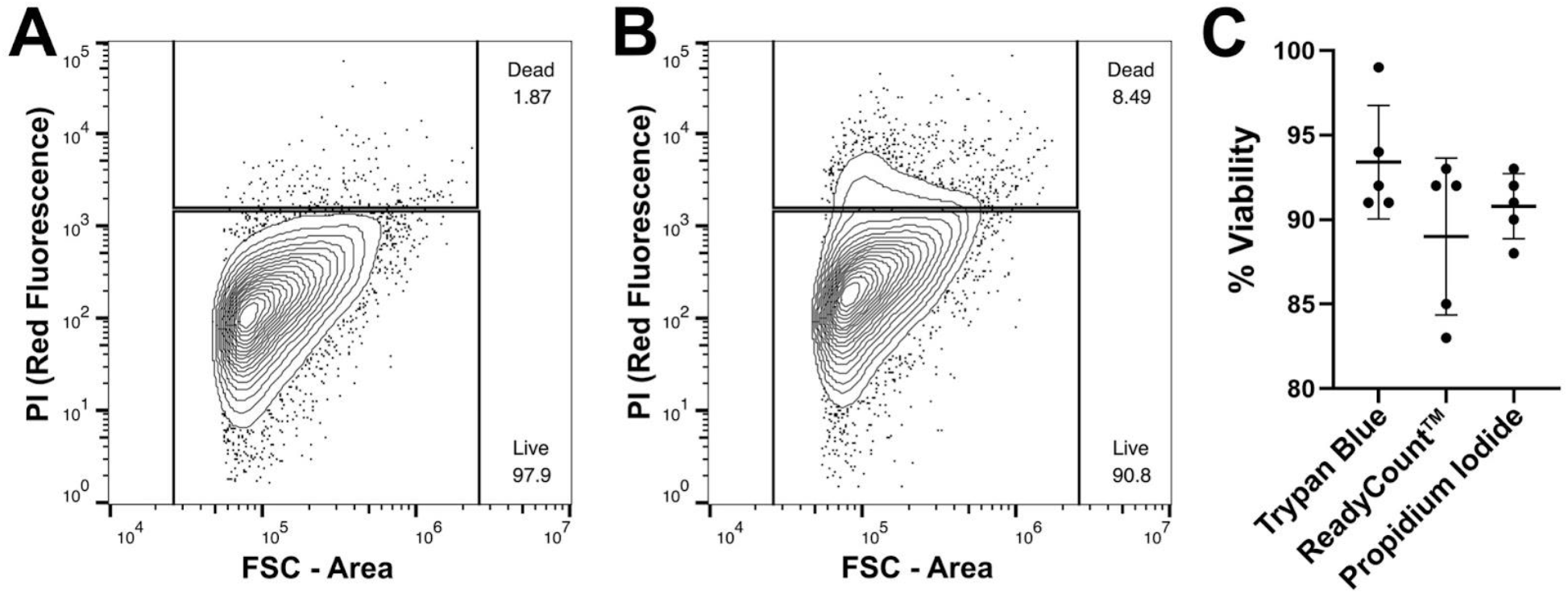
Assessment of *Mnemiopsis* cell preparation viability using DNA staining. A) FACS plot of cell suspension that has not been stained with propidium iodide (PI). There are few events detected in the red fluorescent channel. B) FACS plot of a *Mnemiopsis* cell suspension that has been stained with PI. Increased red fluorescence signal indicates the labeling of apoptotic or necrotic cells, comprising 8.49% of the sample. C) Comparisons of *Mnemiopsis* cell suspension viability using Trypan Blue exclusion assays, ReadyCount Green/Red Viability Stain, or PI. There was no significant difference in cell viability quantification between stains.

Replicates from the same cell suspensions were tested for all three cell viability dyes to compare consistency. We found no statistically significant difference (*p* = 0.18; Table 1; Figure 2C). Thus either PI, Trypan Blue, or ReadyCount Green/Red stains can be used to quantitatively and accurately assess viability of *Mnemiopsis* cell preparations.

### 3.2 Analyzing relative cell size and granularity

Measurements of FSC/SSC can facilitate broad identification of cell morphologies within a sample by separating cells based on variation in relative cell size, intracellular granularity and/or membrane complexity (Rico et al., 2021). To initially examine general morphological characteristics within heterogeneous cell populations prepared from whole *Mnemiopsis*, we measured FSC/SSC of unstained cell preparations. Representative results from FSC/SSC analyses using a Sony SH800 identified five broad cell clusters (Figure 3; Supplemental Figure 2). Following exclusion of doublet and multiplet events representing cell aggregates (Figure 1B), we analyzed and gated on populations of events representing variable FSC and SSC values. Microscopy on sorted live *Mnemiopsis* cells revealed expected correlations between gate selection, relative cell size, and intracellular granule complexity. For example, cells with the lowest values for FSC and SSC isolated from Gate A are relatively small and have few or no visible granules. As FSC-A and SSC-A values increase, cells increase in size and/or morphological complexity. Gates A and B contain small, agranular or semi-granular cells that were highly abundant, while Gates C and E capture large highly granular cells that were the least abundant (Figure 3; Supplemental Figure 2).

**Figure 3:**
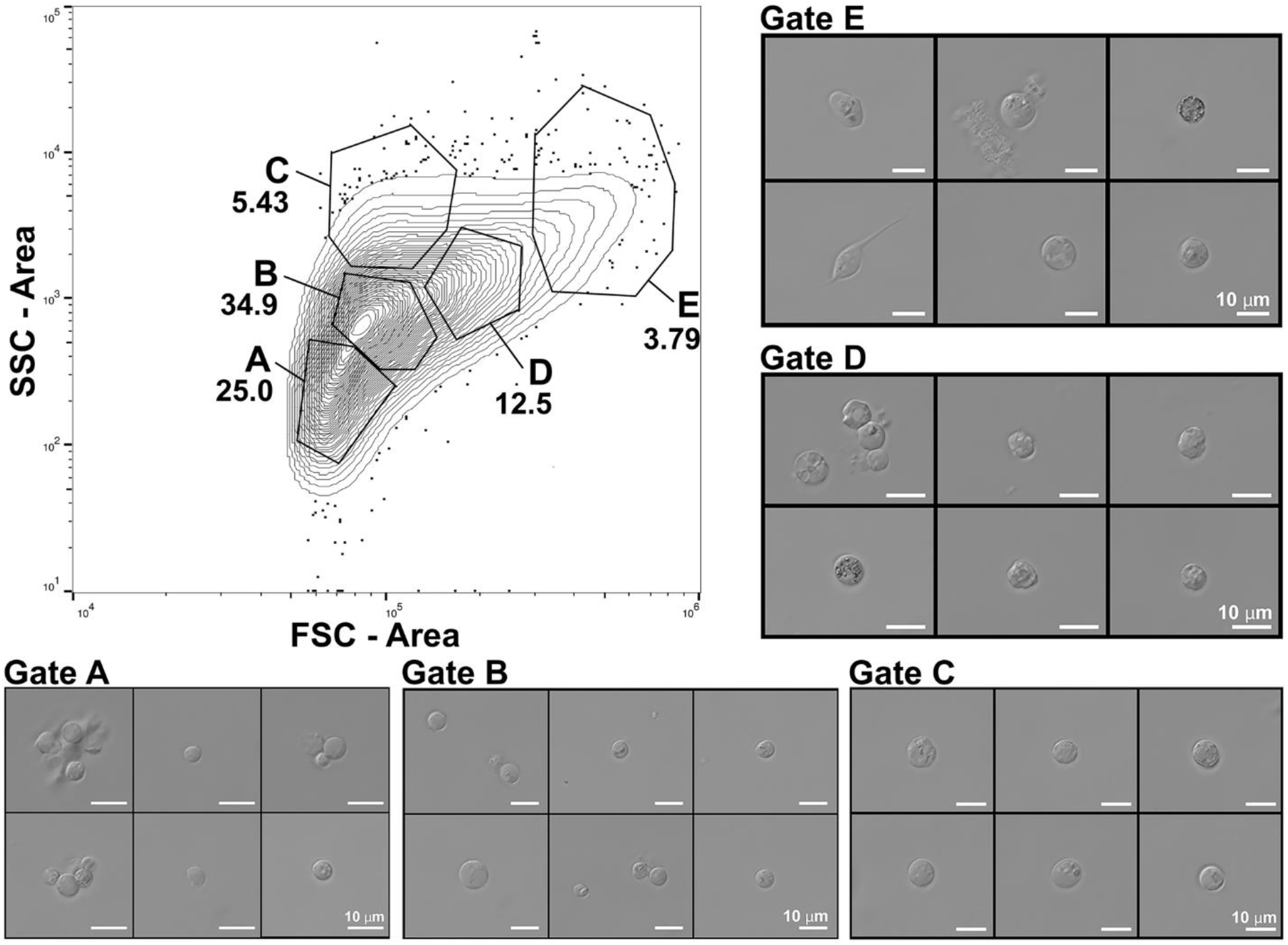
Assessment of general morphological characteristics within heterogeneous unstained cell populations prepared from whole *Mnemiopsis*. Representative contour plot (2%) of FSC-A/SSC-A profiles of a cell suspension with outliers shown. FSC/SSC analyses identified five broad cell clusters. DIC microscopy on sorted *Mnemiopsis* cells shows expected correlations between relative cell size, intracellular complexity and gate selection. Cells with the lowest values for FSC and SSC isolated from Gate A are relatively small with few or no visible granules. As FSC-A and SSC-A values increase, cells increase in size and morphological complexity. Gates C, D and E capture larger, highly granular cells.

### 3.3 Determining cell cycle stages in *Mnemiopsis* cells by FACS

Nuclear DNA in *Mnemiopsis* cells was stained with Vybrant DyeCycle Green to assess cell cycle state distribution. FACS plots comparing Vybrant Green-positive cells demonstrate that cells of variable sizes have the similar DNA content (Figure 4A). This result was expected for mixed cell populations containing an array of cell types when querying by FSC-A (refer to Figure 3; Qiu et al., 2013; Kim and Sedersom, 2015). A threshold gate on the FITC axis can be used to remove background fluorescent signal representing dead and dying/apoptotic cells containing less than 2N DNA content (Vignon et al., 2013).

**Figure 4:**
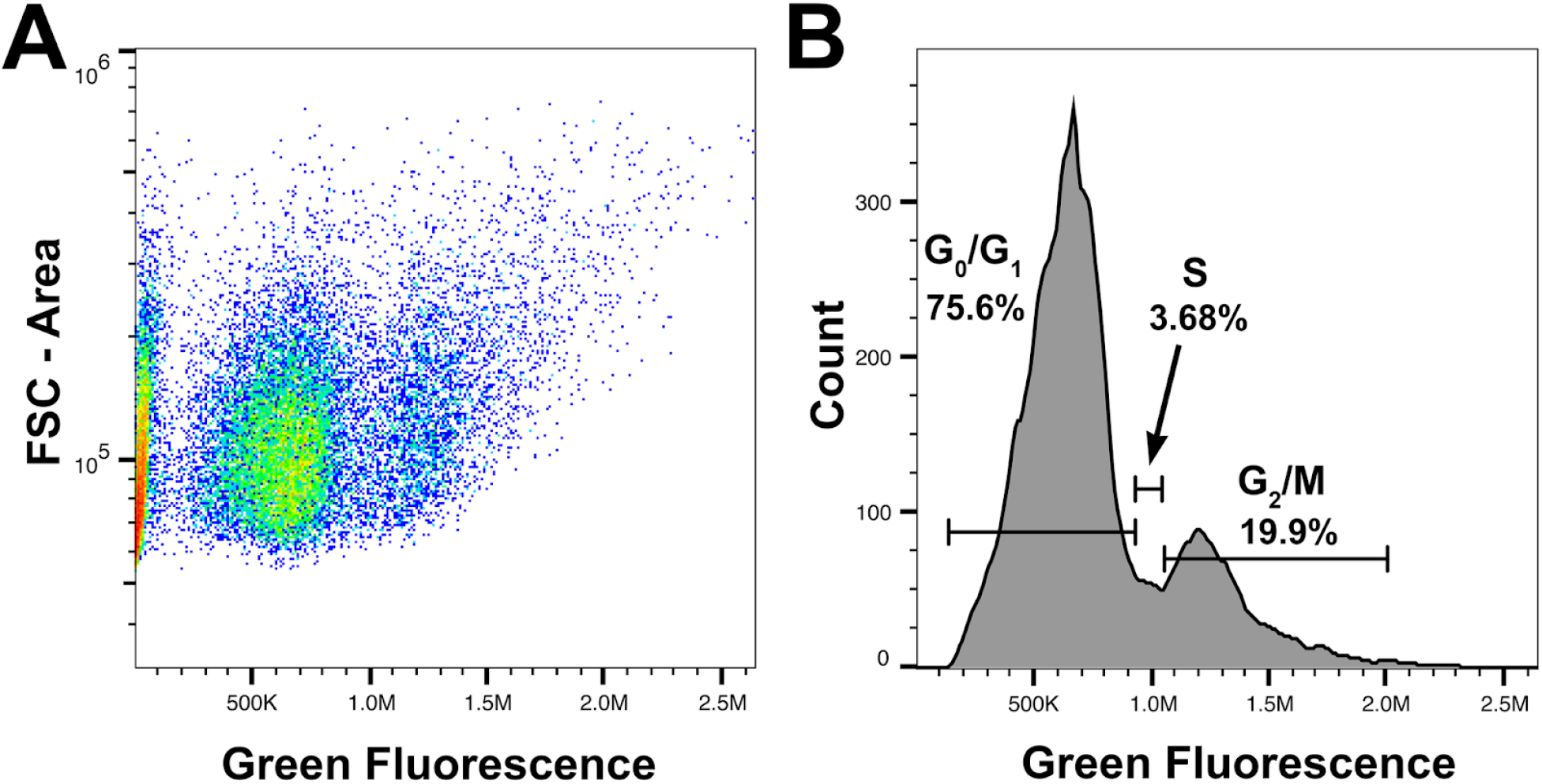
Analysis of cell cycle state distribution using Vybrant DyeCycle. A) Representative FACS plot demonstrating labeling of Vybrant DyeCycle in *Mnemiopsis* cells. Two clusters of cells are visible based on green fluorescence intensity. The FSC-A axis shows that cells of variable sizes have similar DNA content. B) Histogram of green fluorescence reveals two distinct peaks of fluorescence representing cells in G0/G1, S, or G2/M. Events below 10^5^ are background fluorescent signal or represent dead and dying/apoptotic cells that contain less than 2N DNA content.

We observed three distinct cell populations by DNA labeling: cells in G0/G1, S phase, or G2/M (Figure 4B; Vignon et al., 2013; Kim and Sedersom, 2015). Cells in G0 are quiescent and have a genome content of 2N, while cells in G1 also have a DNA content of 2N and are preparing to initiate DNA replication in addition to performing other normal cellular functions. Cells in S and G2/M (4N) are in the process of actively replicating DNA and dividing, and have a DNA content of 4N. This increase in DNA content can be seen as a second fluorescent signal peak across replicate samples (Figure 4B; Supplemental Fig. 3).

### 3.4 Identification and isolation of phagocytic cells

Phagocytosis is a fundamental cell behavior central to many metazoan cell processes, including nutrient uptake and immune response (Hartenstein and Martinez, 2019). Cells competent for phagocytosis can be readily identified using FACS by selecting cells that have internalized fluorescent particles (Lehmann et al., 2000). To assess whether FACS could be applied to target, isolate and collect *Mnemiopsis* phagocytes, we incubated heterogeneous cell suspensions with pHrodo *E. coli* BioParticles along with MCM alone as control cell suspensions. We analyzed samples on both Sony SH800 and BS FACSAria Fusion flow cytometry systems to compare fluorescent and FSC/SSC profiles (Figure 5). Control samples show a normal FSC/SSC profile (Figure 5A, 5F, compare to Figure 1C) and low levels of fluorescence (Figure 5B, 5G). The FSC/SSC profile of pHrodo incubated samples show an increase in SSC, indicative that intracellular granularity has increased in cells that have phagocytosed bacteria (Figure 5C, 5H). We also observed a shift in fluorescent signal in approximately 9-15% of cells across replicates, signifying that these cells had sequestered bacteria into low pH vesicles, activating fluorescence (pHrodo-positive cells; Figure 5C-D, 5H-I; Supplemental Figure 4).

**Figure 5:**
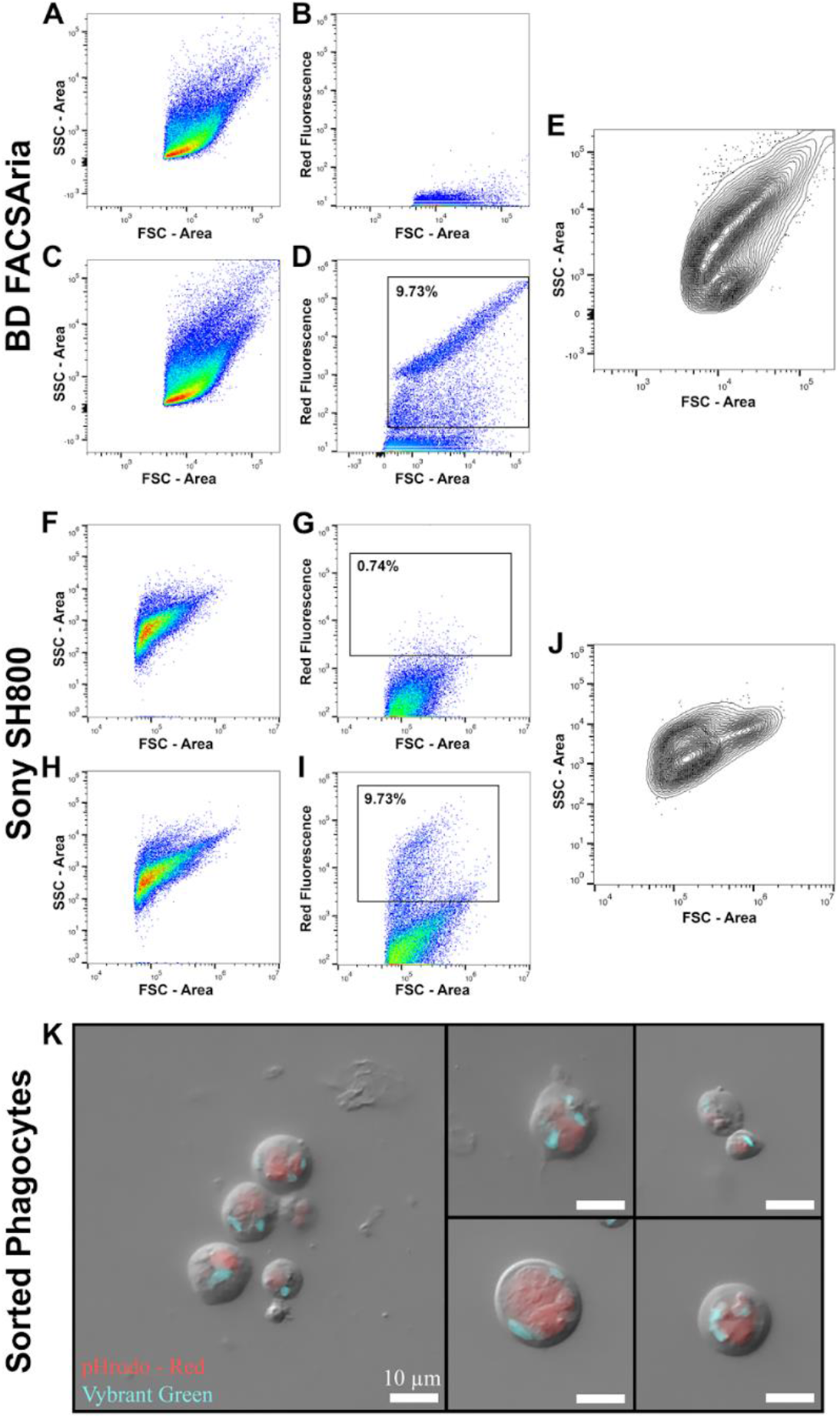
Analysis of phagocytic ability of *Mnemiopsis* cells by FACS. A-E) Data collection from BD FACSAria™ Fusion Flow Cytometer. F-J) Data collection from a Sony SH800 Cell Sorter. A,F) FSC-A/SSC-A dot plot of control *Mnemiopsis* cell suspensions not incubated with phRodo *E. coli*. B,G) Control sample showing no red fluorescence. C,H) FSC-A/SSC-A dot plot of *Mnemiopsis* cell suspensions that have been incubated with pHrodo *E. coli*. A shift in events with higher SSC-A are observed, indicating that some cells have ingested bacteria thereby increasing intracellular granularity. D,I) Positive shift in red fluorescent signal indicating that some cells have phagocytosed pHrodo *E. coli* and trafficked the bacteria to acidic vesicles, activating fluorescence. FSC-A shows that phagocytic cells are diverse in size. E,J) Back-gating on selected fluorescence-positive cells shows that cells of diverse sizes and intracellular complexities phagocytose bacteria. K) Combined DIC and fluorescent microscopy on sorted phagocytic *Mnemiopsis* cells. A variety of cell types have phagocytosed the fluorescent *E. coli*. Red: pHrodo *E. coli* bioparticles. Blue: nuclei. Scale bar 10 um in all panels.

We sorted pHrodo-positive cells and labeled cell nuclei with Vibrant Green to better visualize and identify intact phagocytic cells. We recovered an array of cells of varying sizes and intracellular complexities, including cells that look similar to previously reported round digestive cells (Figure 5E, 5J-K; Presnell et al., 2016; Vandepas et al., 2017; Traylor-Knowles et al., 2019). Intriguingly, some phagocytes also display multiple processes (Figure 5K). Stellate cells with phagocytic capability have been identified in *Mnemiopsis* and may have immune function (Traylor-Knowles et al., 2019). We show here that a range of *Mnemiopsis* phagocytic cells can be identified, isolated and collected using FACS.

## 4 Discussion

The standardized methods presented here allow for the preparation of ctenophore cells from whole animals for reproducible flow cytometry analyses. We demonstrate techniques critical for filtering debris and removing cell aggregates from heterogenous ctenophore cell suspensions to efficiently and accurately identify intact single cells. Our results demonstrate that applying selective gating facilitates the isolation of targeted populations of live, unstained *Mnemiopsis* cells over a range of sizes and intracellular or morphological complexities. We also detail DNA staining parameters for determining cell viability, as well as analysis of cell cycle state. Many species of ctenophores, including *Mnemiopsis*, are capable of rapid regeneration and wound healing of damaged body parts (Henry and Martindale, 2000; Edgar et al., 2021). The ability to analyze cell proliferation by FACS in ctenophores will be a useful tool in studying the remarkable regenerative properties of this clade.

We performed phagocytosis assays as an explicit functional approach for using FACS to analyze and isolate *Mnemiopsis* cells. We identified functionally phagocytic cell populations in *Mnemiopsis* and show via FSC/SSC and microscopy that cells of diverse size and morphology are capable of ingesting and sequestering bacteria in phagosomes. Additional FACS experiments on *Mnemiopsis* phagocytic cells such as co-labeling with reactive oxygen species (ROS) and other identifiers of cellular mechanisms will facilitate further characterization of phagocytic cell types (Rosental et al., 2017).

These flow cytometry methods have significant implications for the study of ctenophore cell biology. The ability to isolate specific ctenophore cell types by FACS will enable a wide range of downstream applications such as gene expression studies, epigenetic profiling, and immune response assays. In the future, optimization of additional fluorescent reagents to query ctenophore cells - such as tagged antibodies - will enable enhanced isolation techniques, as well as the identification of additional specific ctenophore cell types. These applications will improve our understanding of specific ctenophore cell types, behaviors, and cellular processes, providing insight into both the conservation and divergence of cellular processes across Metazoa.

## Author Contributions

Conceptualization, L.E.V., W.E.B., N.T.K., A.L.H. Methodology, L.E.V., E.G., A.L.H.

Formal Analysis, L.E.V., E.G., A.C.D., W.E.B.

Investigation, L.E.V., A.C.D., E.G., A.B.T., W.E.B.

Data Curation, L.E.V., W.E.B. Visualization, L.E.V., A.C.D., E.G., W.E.B.

Writing - Original Draft, A.C.D, L.E.V., W.E.B. Writing - Review and Editing, All authors Resources, N.T.-K., W.E.B., A.L.H.

Supervision, L.E.V., W.E.B, N.T.-K. Funding Acquisition, N.T.-K., W.E.B.

## Acknowledgements

The authors are grateful to the Benaroya Research Institute’s Flow Cytometry Core for technical support.

## FIGURES

**Supp. Figure 1:**
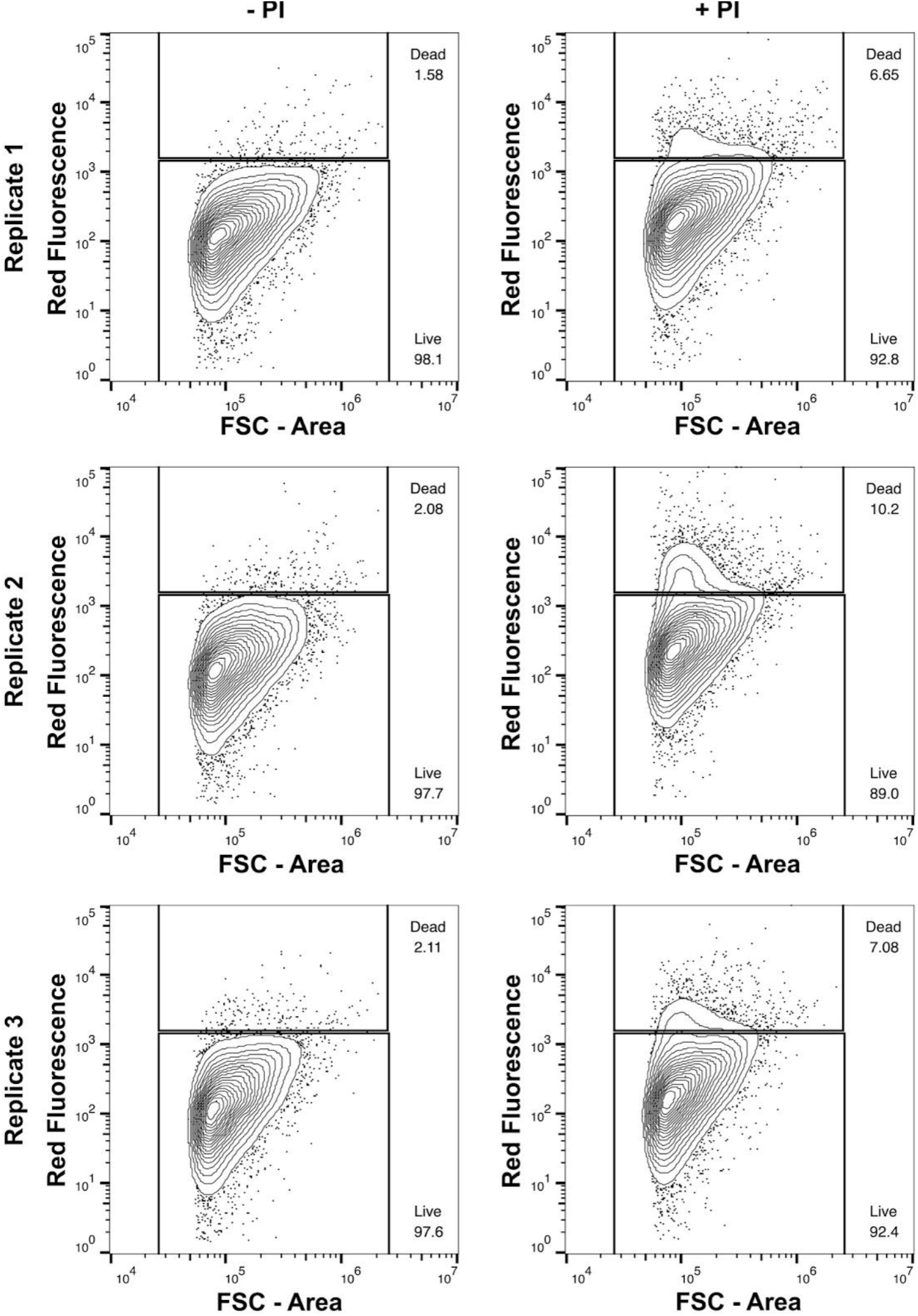
Propidium iodide live/dead assay on *Mnemiopsis* cell suspensions across three biological replicates.

**Supp. Figure 2:**
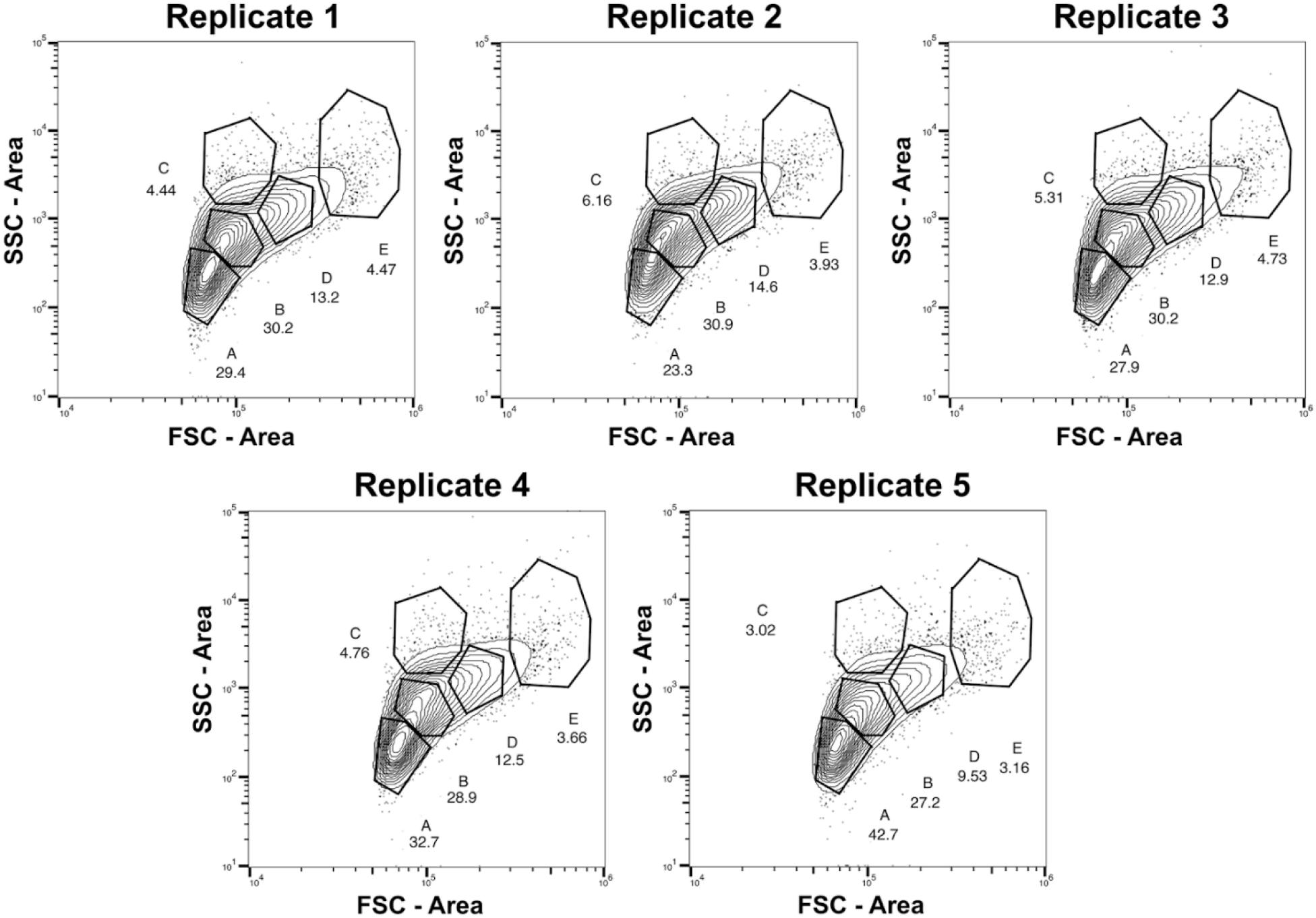
Flow cytometry profiles of unstained *Mnemiopsis* cell suspensions across five biological replicates. Contour plot (2%) showing FSC-A/SSC-A of *Mnemiopsis* cell suspensions with gating strategies and corresponding frequencies of each population.

**Supp. Figure 3:**
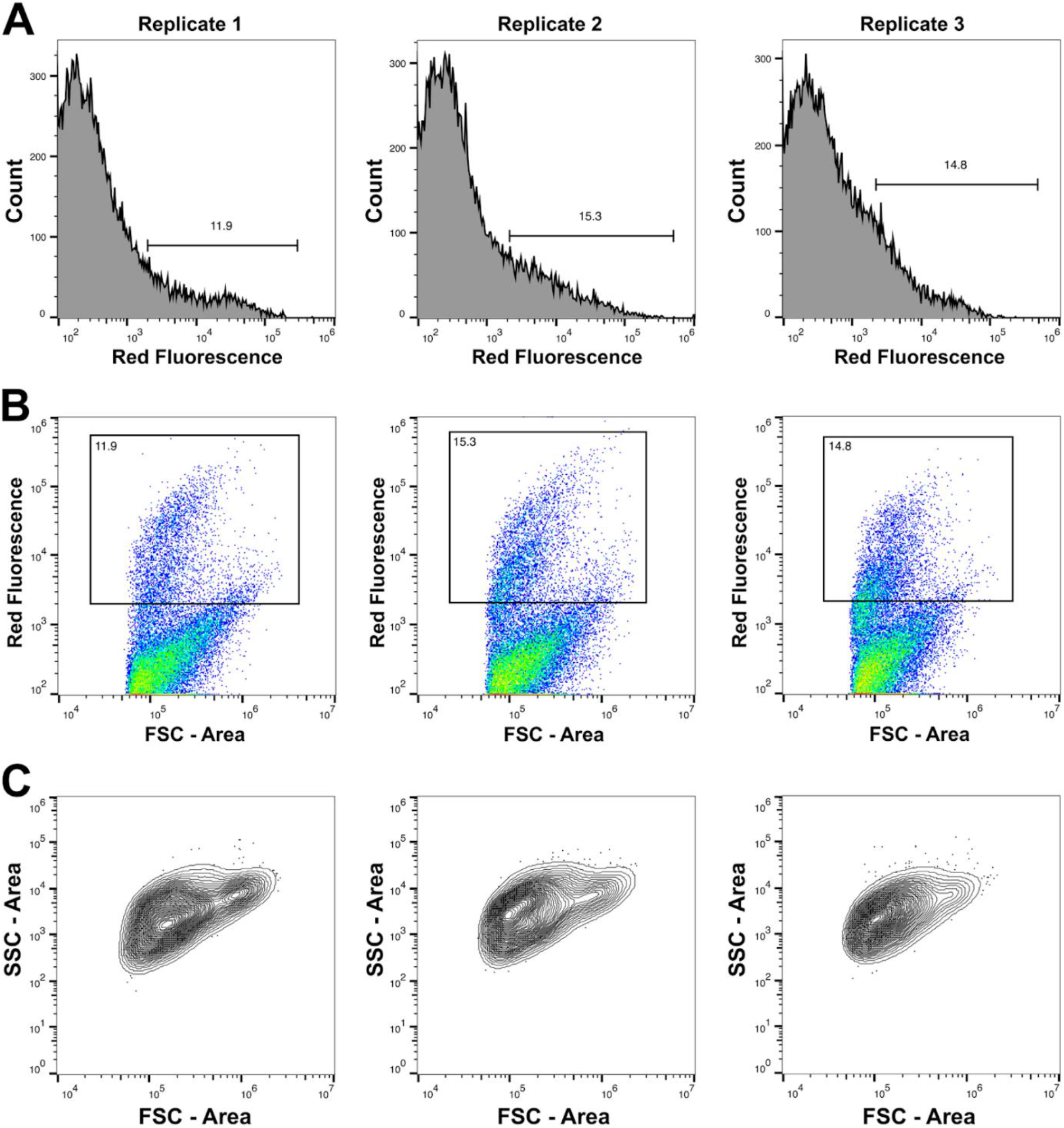
Phagocytosis assays using pHrodo *E. coli* Bioparticles in *Mnemiopsis* cell suspensions across four biological replicates. A) Histograms showing distribution of fluorescent signals and gating on pHrodo-positive cells across four biological replicates. B) Flow cytometry profiles of *Mnemiopsis* cell suspensions showing that cells of diverse sizes, as shown by FSC-A, have phagocytosed *E. coli*. C) Back-gating on pHrodo-positive cells shows that cells displaying a variety of sizes and complexities are capable of phagocytosing bacteria.

## References

Barteneva NS, Fasler-Kan E, Vorobjev IA. (2012) Imaging flow cytometry: coping with heterogeneity in biological systems. J Histochem Cytochem. 60(10), pp. 723–733. doi: 10.1369/0022155412453052.

Babonis, L.S., DeBiasse, M.B., Francis, W.R., Christianson, L.M., Moss, A.G., Haddock, S.H., Martindale, M.Q. and Ryan, J.F., 2018. Integrating embryonic development and evolutionary history to characterize tentacle-specific cell types in a ctenophore. Molecular Biology and Evolution, 35(12), pp.2940–2956.

Bradford, J.A., Whitney, P., Huang, T., Pinson, P., Cheung, C.-Y., Yue, S. and Godfrey, W.L. (2006). Novel Vybrant® DyeCycle TM Stains Provide Cell Cycle Analysis in Live Cells Using Flow Cytometry with Violet, Blue, and Green Excitation.. Blood, [online] 108(11), p. 4234. doi:10.1182/blood.V108.11.4234.4234.

Burkhardt, P., Colgren, J., Medhus, A., Digel, L., Naumann, B., Soto-Angel, J.J., Nordmann, E.L., Sachkova, M.Y. and Kittelmann, M., 2023. Syncytial nerve net in a ctenophore adds insights on the evolution of nervous systems. Science, 380(6642), pp.293–297.

Cheng, S.F., Cai, X., Deng, D. and Wang, W., 2018. Classification of Hemocytes from Four Crustaceans and Cross-Reactivity of Their Antisera. Journal of Shellfish Research, 37(1), pp.159–171.

Choi, D.L., Lee, N.S., Kim, M.S., Seo, J.S., Park, M.A., Kim, J.W. and Hwang, J.Y., 2010. Flow cytometry analysis of softness syndrome effects on hemocytes of the tunicate Halocynthia roretzi. Aquaculture, 309(1-4), pp. 25–30.

Dayraud, C., Alié, A., Jager, M., Chang, P., Le Guyader, H., Manuel, M. and Quéinnec, E. (2012). Independent specialisation of myosin II paralogues in muscle vs. non-muscle functions during early animal evolution: a ctenophore perspective. BMC Evolutionary Biology, 12(1). doi:10.1186/1471-2148-12-107.

de la Cruz, A.F.A. and Edgar, B.A., 2008. Flow cytometric analysis of Drosophila cells. Drosophila: Methods and Protocols, pp. 373-389.

Dieter, A.C., Vandepas, L.E., Browne, W.E. (2022). Isolation and Maintenance of In Vitro Cell Cultures from the Ctenophore Mnemiopsis leidyi. Methods in Molecular Biology, 2450., pp. 347–358. https://doi.org/10.1007/978-1-0716-2172-1_18

Dunn, C.W., Leys, S.P. and Haddock, S.H., 2015. The hidden biology of sponges and ctenophores. Trends in ecology & evolution, 30(5), pp.282–291.

Edgar, A., Mitchell, D.G. and Martindale, M.Q., 2021. Whole-body regeneration in the lobate ctenophore Mnemiopsis leidyi. Genes, 12(6), p. 867.

Ellis, R.P., Parry, H., Spicer, J.I., Hutchinson, T.H., Pipe, R.K. and Widdicombe, S. (2011). Immunological function in marine invertebrates: Responses to environmental perturbation. Fish & Shellfish Immunology, 30(6), pp. 1209–1222. doi:10.1016/j.fsi.2011.03.017.

Fulwyler, M.J., 1965. Electronic separation of biological cells by volume. Science, 150(3698), pp. 910–911.

Hartenstein V, Martinez P. Phagocytosis in cellular defense and nutrition: a food-centered approach to the evolution of macrophages. Cell Tissue Res. 2019 Sep;377(3):527–547. doi: 10.1007/s00441-019-03096-6. Epub 2019 Sep 4. PMID: 31485720; PMCID: PMC6750737.

Henry, J.Q. and Martindale, M.Q., 2000. Regulation and regeneration in the ctenophore Mnemiopsis leidyi. Developmental biology, 227(2), pp. 720–733.

Herzenberg, L.A., Parks, D., Sahaf, B., Perez, O., Roederer, M. and Herzenberg, L.A. (2002). The history and future of the fluorescence activated cell sorter and flow cytometry: a view from Stanford. Clinical Chemistry, [online] 48(10), pp. 1819–1827.

Hernandez-Nicaise ML (1991). Ctenophora. In: Harrison FW (ed) Microscopic anatomy of the invertebrates, vol 2. Wiley-Liss, New York, pp 359–418

Jager, M., Chiori, R., Alié, A., Dayraud, C., Quéinnec, E. and Manuel, M. (2010). New insights on ctenophore neural anatomy: Immunofluorescence study in Pleurobrachia pileus (Müller, 1776). Journal of Experimental Zoology Part B: Molecular and Developmental Evolution, 316B(3), pp. 171–187. doi:10.1002/jez.b.21386.

Johnson, S., Nguyen, V., & Coder, D. (2013). Assessment of cell viability. Current Protocols in Cytometry, 64(1), 9.2.1–9.2.26.

Jokura, K., Sato, Y., Shiba, K. and Inaba, K., 2022. Two distinct compartments of a ctenophore comb plate provide structural and functional integrity for the motility of giant multicilia. Current Biology, 32(23), pp. 5144–5152.

Julius, M.H., Masuda, T. and Herzenberg, L.A. (1972). Demonstration That Antigen-Binding Cells Are Precursors of Antibody-Producing Cells After Purification with a Fluorescence-associated Cell Sorter. Proceedings of the National Academy of Sciences, 69(7), pp. 1934–1938. doi:10.1073/pnas.69.7.1934.

Kim, K.H. and Sederstrom, J.M., 2015. Assaying cell cycle status using flow cytometry. Current protocols in molecular biology, 111(1), pp. 28–6.

Kissing S, Saftig P, Haas A. (2018) Vacuolar ATPase in phago(lyso)some biology. Int J Med Microbiol. 308(1), pp. 58–67. doi: 10.1016/j.ijmm.2017.08.007..

Kron, P., Suda, J. and Husband, B.C. (2007). Applications of Flow Cytometry to Evolutionary and Population Biology. Annual Review of Ecology, Evolution, and Systematics, 38(1), pp. 847–876. doi:10.1146/annurev.ecolsys.38.091206.095504.

Kummrow, A., Frankowski, M., Bock, N., Werner, C., Dziekan, T. and Neukammer, J., 2013. Quantitative assessment of cell viability based on flow cytometry and microscopy. Cytometry Part A, 83(2), pp. 197–204.

Lehmann, A.K., Sørnes, S. and Halstensen, A., 2000. Phagocytosis: measurement by flow cytometry. Journal of immunological methods, 243(1-2), pp.229-242.

Li, Y., Shen, X.X., Evans, B., Dunn, C.W. and Rokas, A., 2021. Rooting the animal tree of life. Molecular Biology and Evolution, 38(10), pp. 4322–4333.

Marringa, W.J., Krueger, M.J., Burritt, N.L. and Burritt, J.B. (2014). Honey Bee Hemocyte Profiling by Flow Cytometry. PLoS ONE, 9(10), p. e108486. doi:10.1371/journal.pone.0108486.

McKinnon, Katherine M. (2018) “Flow Cytometry: An Overview.” Current protocols in immunology 120:5.1.1-5.1.11. 21 doi:10.1002/cpim.40

Moroz, L.L., Kocot, K.M., Citarella, M.R., Dosung, S., Norekian, T.P., Povolotskaya, I.S., et al. (2014). The ctenophore genome and the evolutionary origins of neural systems. Nature, 510(7503), pp. 109–114. doi:10.1038/nature13400.

Park, K.-I., Donaghy, L., Kang, H.-S., Hong, H.-K., Kim, Y.-O. and Choi, K.-S. (2012). Assessment of immune parameters of manila clam Ruditapes philippinarum in different physiological conditions using flow cytometry. Ocean Science Journal, 47(1), pp. 19–26. doi:10.1007/s12601-012-0002-x.

Presnell, Jason S., Vandepas Lauren E., Warren Kaitlyn J., Swalla Billie J., Amemiya Chris T. and Browne William E. (2016). The Presence of a Functionally Tripartite Through-Gut in Ctenophora Has Implications for Metazoan Character Trait Evolution. Current Biology, 26(20), pp. 2814–2820. doi:10.1016/j.cub.2016.08.019.

Presnell, J. S., Bubel, M., Knowles, T., Patry, W., Browne, W. E.. (2022) Multigenerational laboratory culture of pelagic ctenophores and CRISPR/Cas9 genome editing in the lobate, Mnemiopsis leidyi. Nature Protocols 17, 1868–1900.

Qiu, L., Liu, M. and Pan, K., 2013. A triple staining method for accurate cell cycle analysis using multiparameter flow cytometry. Molecules, 18(12), pp. 15412–15421.

Riccardi, C. and Nicoletti, I., 2006. Analysis of apoptosis by propidium iodide staining and flow cytometry. Nature protocols, 1(3), pp. 1458–1461.

Rico, L.G., Salvia, R., Ward, M.D., Bradford, J.A. and Petriz, J., 2021. Flow-cytometry-based protocols for human blood/marrow immunophenotyping with minimal sample perturbation. STAR protocols, 2(4), p. 100883.

Rosental, B., Kowarsky, M.A., Corey, D.M., Ishizuka, K.J., Palmeri, K.J., Chen, S.Y., Sinha, R., Voskoboynik, A. and Weissman, I.L., 2016. Finding the evolutionary precursor of vertebrate hematopoietic lineage: Functional and molecular characterization of B. schlosseri immune system. The Journal of Immunology, 196(1_Supplement), pp.216-2.

Rosental, B., Kozhekbaeva, Z., Fernhoff, N., Tsai, J.M. and Traylor-Knowles, N., 2017. Coral cell separation and isolation by fluorescence-activated cell sorting (FACS). BMC Cell Biology, 18, pp. 1–12.

Schippers, K. J., Martens, D. E., Pomponi, S. A., & Wijffels, R. H. (2011). Cell cycle analysis of primary sponge cell cultures. In vitro cellular & developmental biology. Animal, 47(4), pp. 302–311. doi.org:10.1007/s11626-011-9391-x.

Sebé-Pedrós, A., Chomsky, E., Pang, K., Lara-Astiaso, D., Gaiti, F., Mukamel, Z., Amit, I., Hejnol, A., Degnan, B.M. and Tanay, A. (2018). Early metazoan cell type diversity and the evolution of multicellular gene regulation. Nature Ecology & Evolution, [online] 2(7), pp. 1176–1188. doi:10.1038/s41559-018-0575-6.

Siebert, S., Farrell, J.A., Cazet, J.F., Abeykoon, Y., Primack, A.S., Schnitzler, C.E. and Juliano, C.E., 2019. Stem cell differentiation trajectories in Hydra resolved at single-cell resolution. Science, 365(6451), p. eaav9314.

Snow, C. (2004). Flow cytometer electronics. Cytometry 57A(2): pp. 63-69. doi.org/10.1002/cyto.a.10120

Snyder, G. A., Browne, W. E., Traylor-Knowles, N., Rosental, B. (2020). Fluorescence-Activated Cell Sorting for the Isolation of Scleractinian Cell Populations. Journal of Visualized Experiments (159), 60446, doi:10.3791/60446

Tamm, S. and Tamm, S. (1991). Actin pegs and ultrastructure of presumed sensory receptors of Beroe (Ctenophora). Cell and Tissue Research, 264(1), pp. 151–159. doi:10.1007/bf00305733.

Traylor-Knowles, N., Vandepas, L.E. and Browne, W.E. (2019). Still Enigmatic: Innate Immunity in the Ctenophore Mnemiopsis leidyi. Integrative and Comparative Biology, 59(4), pp. 811–818. doi:10.1093/icb/icz116.

Vandepas, L.E., Warren, K.J., Amemiya, C.T. and Browne, W.E. (2017). Establishing and maintaining primary cell cultures derived from the ctenophore Mnemiopsis leidyi. The Journal of Experimental Biology, 220(7), pp. 1197–1201. doi:10.1242/jeb.152371.

Vignon, C., Debeissat, C., Georget, M.T., Bouscary, D., Gyan, E., Rosset, P. and Herault, O., 2013. Flow cytometric quantification of all phases of the cell cycle and apoptosis in a two-color fluorescence plot. PloS one, 8(7), p. e68425.

